# Phosphoproteomic mapping reveals distinct signaling actions and activation of protein synthesis and muscle hypertrophy by Isthmin-1

**DOI:** 10.1101/2022.05.19.492758

**Authors:** Meng Zhao, Niels Banhos Danneskiold-Samsøe, Livia Ulicna, Quennie Nguyen, Laetitia Voilquin, David E. Lee, James P. White, Zewen Jiang, Nickeisha Cuthbert, Shrika Paramasivam, Ewa Bielczyk-Maczynska, Capucine Van Rechem, Katrin J. Svensson

## Abstract

The secreted protein Isthmin-1 (Ism1) mitigates diabetes by increasing adipocyte and skeletal muscle glucose uptake by activating the PI3K-Akt pathway. However, while both Ism1 and insulin converge on these common targets, Ism1 has distinct cellular actions suggesting divergence in downstream intracellular signaling pathways. To understand the biological complexity of Ism1 signaling, we performed phosphoproteomic analysis after acute exposure, revealing overlapping and distinct pathways of Ism1 and insulin. We identify a 53 % overlap between Ism1 and insulin signaling and Ism1-mediated phosphoproteome-wide alterations in ∼ 450 proteins that are not shared with insulin. Interestingly, we find several unknown phosphorylation sites on proteins related to protein translation, mTOR pathway and, unexpectedly, muscle function in the Ism1 signaling network. Physiologically, Ism1 ablation in mice results in altered proteostasis, including lower muscle protein levels under fed and fasted conditions, reduced amino acid incorporation into proteins, and reduced phosphorylation of the key protein synthesis effectors Akt and downstream mTORC1 targets. As metabolic disorders such as diabetes are associated with accelerated loss of skeletal muscle protein content, these studies define a non-canonical mechanism by which this anti-diabetic circulating protein controls muscle biology.

## Introduction

Hormonal signaling through protein phosphorylation is one of the most important post-translational modifications allowing for rapid changes in cellular metabolic states (Humphrey et al., 2015). Metabolic stressors such as diabetes or fasting can lead to pronounced physiologic and cellular adaptations in protein regulation (Powers et al., 2009). Consequently, pathologic conditions can result in muscle atrophy, a net loss of muscle mass, which is highly associated with morbidity (Cohen et al., 2015; Jackman and Kandarian, 2004). Muscle strength is reduced in individuals with insulin resistance and type 2 diabetes (Andersen et al., 2004; Park et al., 2007), and muscle weakness is even a diagnostic predictor of diabetes (Peterson et al., 2016). The mechanisms underlying this association have remained elusive, but it is plausible that factors derived from adipose tissue can contribute to muscle proteostasis. Identifying such molecular triggers of muscle metabolism could facilitate our efforts to develop pharmacological agents that improve muscle function and systemic metabolic health.

Skeletal muscle is the most abundant tissue in humans, representing up to half of the total mass in normal-weight individuals (Janssen et al., 2000). As a major organ for glycogen storage and insulin-mediated glucose uptake, skeletal muscle controls whole-body energy expenditure and nutrient homeostasis (Deshmukh, 2016; Shulman et al., 1990). Importantly, skeletal muscle also acts as a protein reservoir that is highly responsive to anabolic or catabolic hormonal stimulation, including growth hormone (GH) and insulin-like growth factor-1 (IGF-1), both of which stimulate muscle fiber size (hypertrophy) (Moro et al., 2016; Velloso, 2008). Muscle mass is balanced by pathways controlling protein synthesis and protein degradation. The most well-described anabolic signaling pathway that promotes protein synthesis requires Akt/mTORC1 signaling, which robustly induces muscle hypertrophy upon stimulation by growth factors or amino acids (Bodine et al., 2001; Glass, 2011; Lai et al., 2004). In both flies (Scanga et al., 2000) and mammals (Edinger and Thompson, 2002), the PI3K-Akt pathway controls cell size by increasing protein synthesis at the level of translation initiation. Akt also inhibits the catabolic function of FoxO family members, which upon phosphorylation are no longer able to enter the nucleus and turn on the transcription of atrophy genes, including the E3 ubiquitin ligases MuRF1 and Atrogin-1 (Bodine et al., 2001; Bodine and Baehr, 2014; Gomes et al., 2001; Lecker et al., 1999).

We recently reported that the adipocyte-secreted protein Isthmin-1 (Ism1) improves glucose tolerance and insulin resistance by phosphorylating Akt^S473^, which mediates increased adipose and skeletal muscle glucose uptake (Jiang et al., 2021). ISM1 adipose expression and circulating levels are elevated in mice and humans with obesity (Jiang et al., 2021; Ruiz-Ojeda et al., 2022), suggesting that the expression is under nutrient-sensing regulatory control. Intriguingly, Ism1 administration to mice simultaneously prevents hepatic lipid accumulation while increasing protein synthesis in hepatocytes (Jiang et al., 2021), demonstrating that Ism1 governs an anabolic pathway that is molecularly and functionally distinct from insulin. In the present study, we find that Ism1 induces specific Ism1-regulated phosphoproteome changes enriched for proteins controlling protein translation, mTOR signaling, and muscle function. Furthermore, we show that Ism1 is important to maintain skeletal muscle fiber size under fasting, thereby defining a non-canonical mechanism by which Ism1 controls muscle biology.

## Results

### Phosphoproteomics reveals overlapping and distinct pathways of Ism1 and insulin

The PI3K-Akt pathway is a key pathway of convergence for ligands that activate receptor tyrosine kinases (RTKs), including insulin (Humphrey et al., 2013; Luo et al., 2003; Zhao et al., 2020). We recently identified Ism1 as an adipose-secreted protein that increases glucose uptake into fat and muscle by potently activating PI3K-Akt signaling across a range of mouse and human cell types, but unlike insulin, does not induce *de novo* lipogenesis (Jiang et al., 2021). Therefore, the extent to which the entire Ism1-signaling network overlaps with insulin or whether other signaling nodes are involved remains to be determined. Therefore, to increase our understanding of the signaling divergence and obtain an unbiased, more complete view of the Ism1-induced signaling network, we performed phosphoproteomics in the Ism1 and insulin-responsive 3T3-F442A cells. To characterize the Ism1-induced phospho-signaling profile after acute treatment in cells, we used phosphopeptide enrichment with TiO_2_ followed by LC-MS/MS using Orbitrap Elite (Yue et al., 2015; Zhou et al., 2008) (***Figure 1A***). Cells were starved overnight, followed by a 5-minute treatment with 100 nM recombinant Ism1, or 100 nM insulin. As a negative control, bovine serum albumin (BSA), a secreted protein in the same size range as Ism1, was used at 100 nM. Although pAkt induction was more pronounced by insulin, we observed robust activation of pAkt 5 min post-treatment with Ism1, and therefore selected this time point for our analysis (***Figure 1B***). The proteomic experiments were performed in treatment groups of six biological replicates pooled as duplicate samples for the proteomics analysis. In total, ∼7700 raw MS precursor ions (peptides) were acquired, resulting in the identification of unique phosphopeptides on >5000 proteins (***Figure 1 - figure supplement 1***). Principal component analysis (PCA) demonstrates high reproducibility between biological replicates and distinct separation of the Ism1- and insulin-treated groups compared with each other and the albumin control (***Figure 1C***). Interestingly, we identify overlapping and distinct Ism1 and insulin-specific phosphoproteome-wide alterations upon acute stimulation, with BSA as control. First, there is a 53 % overlap between Ism1 and insulin signaling (***Figure 1D***). For example, insulin induces phosphorylation of 654 phosphosites, out of which 347 phosphosites are also phosphorylated or dephosphorylated by Ism1 (***Figure 1D***).

**Figure 1.**
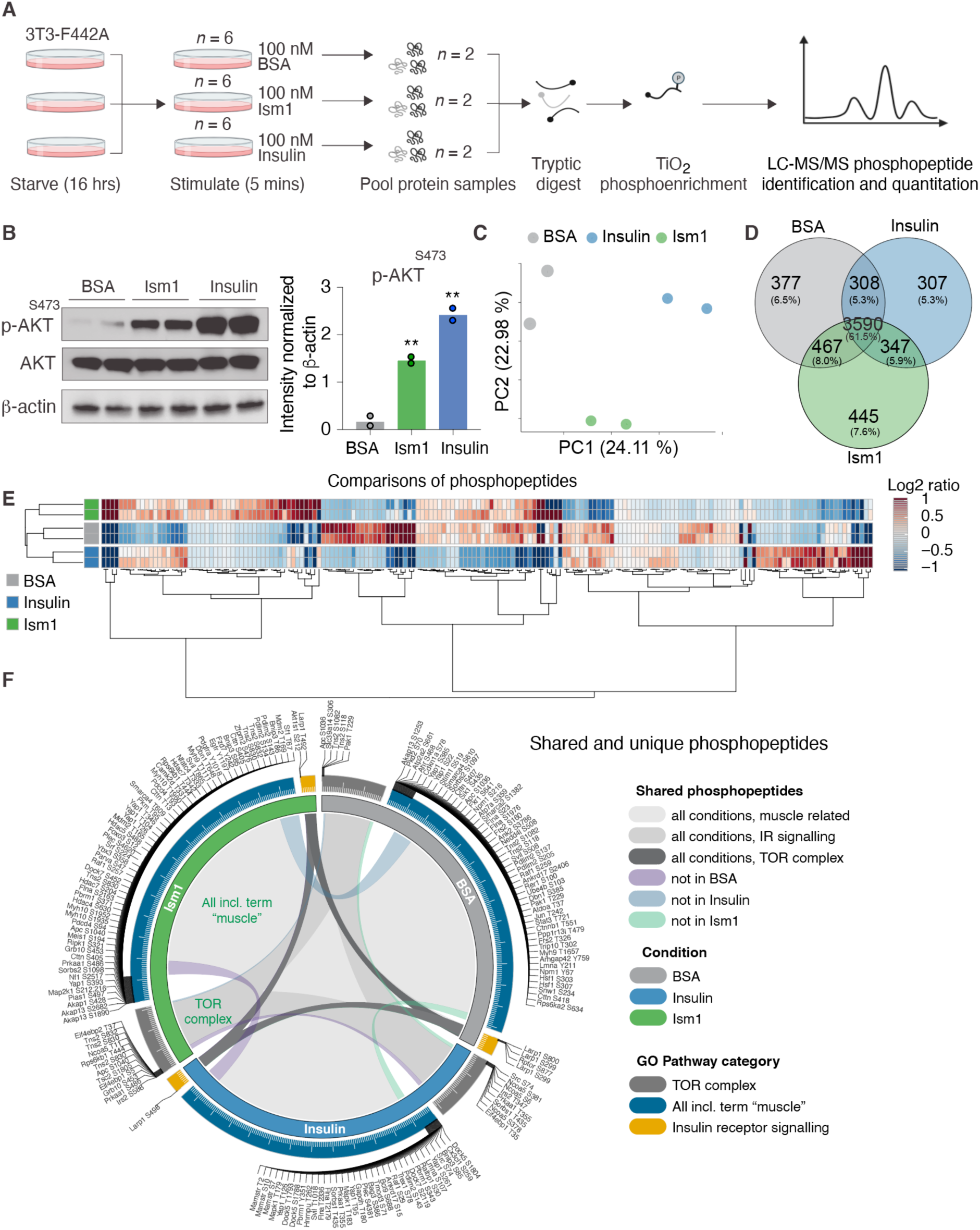
Phosphoproteomics reveals overlapping and distinct pathways of Ism1 and insulin. (A) Experimental design of the untargeted phosphoproteomics analysis. 3T3-F442A cells were serum starved for 16 hours and treated with 100 nM recombinant Ism1 or insulin for 5 minutes (*n* = 6 per group). Proteins were extracted, trypsin digested, and fractionated. Phosphopeptides were enriched using TiO2 chromatography and phosphopeptides were analyzed with LC-MS/MS. (B) Western blot analysis of p-AKT^S473^, total AKT, and β-actin in cells treated with 100 nM bovine serum albumin (BSA), 100 nM Ism1, or 100 nM insulin for 5 minutes (*n* = 2 pooled from *n* = 6 per treatment group). P values are calculated by two-tailed Student’s t-test, *p < 0.05, **p < 0.01, ***p < 0.001.). (C) Protein intensity-based principal component analysis (PCA) of the phosphoproteomic dataset. (D) Venn diagram of the phosphopeptides detected in both replicates after BSA-, Ism1-, or insulin treatment. (E) Heatmap of differentially phosphorylated peptides with 100 lowest significant p-values (adj. p < 0.05) from three comparisons displayed as log2 ratio (BSA vs. Ism1, BSA vs. Insulin, and Insulin vs. Ism1). (F) Distribution diagram of shared and unique phosphosites (detected in at least 1 sample) between treatments for selected GO pathways. Inner links in shades show phosphosites detected in two or more treatment conditions. Gray shows phosphosites detected in all samples; purple shows phosphosites detected in both Ism1 and insulin-treated cells; green shows phosphosites detected in both BSA and insulin-treated cells; blue shows phosphosites detected in both BSA and Ism1 treated cells. The middle ring displays GO pathway/pathway group with ticks indicating the number of phosphopeptides. Large ticks indicate 50 phosphopeptides, and small ticks indicate 5 phosphopeptides. The outer ring displays gene symbols and the phosphosite exclusively detected in each treatment. See also Figure 1—figure supplement 1, and Figure 1—source data 1, 2 and 3.

Remarkably, Ism1 causes changes in the phosphorylation status of 445 proteins compared with BSA that are not shared with insulin and not previously described (***Figure 1D***). Groupwise comparisons between treatments show phosphosites selectively regulated by the specific ligands (adj. p value of < 0.05), many of which have not previously been identified (***Figure 1E***). Based on the notion that Ism1 causes alterations in phosphorylation of a subset of proteins, while another subset is shared with insulin, we next used GO analysis to discern cellular signaling pathways associated with Ism1 or insulin. While Ism1 and insulin share the majority of phosphopeptides for detected genes annotated to these GO terms, some phosphorylated residues could only be identified in one condition (***Figure 1F***). Furthermore, we analyzed to which extent the phosphorylation patterns overlapped between the Ism1 and insulin for genes belonging to the GO terms “GO:0008286, insulin receptor signaling”, “GO:0038201, TOR complex”, and GO terms including the word “muscle” (***Figure 1F***). Interestingly, we find that Ism1 exclusively alters the phosphorylation status of proteins involved in the mTOR complex and muscle (***Figure 1F***). These overlapping and distinct signaling nodes may reflect the signaling networks underlying the cell-specific responses.

### Phospho-specific mapping identifies an Ism1-induced signature consistent with protein translation and muscle function

To interrogate the Ism1-specific signal transduction pathway in more detail, we compared the overlapping phosphopeptides clustered by functional GO pathway groups. Expectedly, insulin induces robust phosphorylation of a subset of proteins, including the insulin receptor (IR). This phosphoproteomic mapping shows that Ism1 does not induce the exact same targets in the insulin pathway as insulin (***Figure 2A***), which is entirely consistent with our previous study using phospho-tyrosine antibodies for the IR (Jiang et al., 2021). For example, only insulin phosphorylates the InsR at Y1175/Y1163 while no significant phosphorylation is induced by Ism1 (***Figure 2B***). Interestingly, Ism1 induces phosphorylation of some of the same proteins as insulin, including insulin receptor substrates Irs1 and Irs2, but with distinct phosphosite patterns. For example, Irs2 was phosphorylated at T347 by insulin but at S588 by Ism1 (***Figure 1F, Figure 2 - figure supplement 1***). Therefore, Ism1 and insulin activate overlapping but distinct pathways, which may account for some of the phenotypic and cell-type specific functions of Ism1. Similar clustering for the mTOR and muscle pathways (***Figure 2C-D***) revealed several proteins regulated only by insulin or only by Ism1. Both ligands induce phosphorylation of tuberin (Tsc2), a negative regulator of mTOR with established roles in the control of protein synthesis (Inoki et al., 2003)(Huang and Manning, 2008). Interestingly, insulin induces Tsc2^S1806^ phosphorylation (***Figure 2E***), while Ism1 induces Tsc2^S1421^ phosphorylation (***Figure 2F***). Additionally, four other phosphosites on Tsc2 are unchanged by Ism1. While the function of these specific phosphosites of Tsc2 are understudied, our data suggest a direct regulation of mTOR activity, potentially functionally distinct from insulin. Furthermore, Ism1 induced phosphorylation of several proteins shown to regulate muscle growth and fiber size, such as Yes1 associated transcriptional regulator (Yap1)^T95^ (Watt et al., 2015) (***Figure 2G***), Heat shock transcription factor 1 (Hsf1)^S236^(Nishizawa et al., 2013) (***Figure 2H***), and FGF-receptor substrate 2 (Frs2)^S327^ (***Figure 2I***) (Chen and Friesel, 2009). Frs2 is also a lipid-anchored adapter protein and downstream mediator of signaling of multiple receptor tyrosine kinases supporting the existence of a distinct Ism1 receptor (Zhou et al., 2009). As protein translation is a critical downstream target of mTOR and the functionally controls muscle growth, we next investigated the regulation of ribosomal targets that catalyze protein synthesis. Interestingly, eukaryotic translation initiation factor 3 subunit A (Eif3), a complex-subunit playing a major role in translation initiation (Ma et al., 2022), was phosphorylated by insulin but not Ism1 (***Figure 2J***), while 60S ribosomal protein L12 (Rpl13) at S77 (***Figure 2K***) and 40S ribosomal protein S13-1 (Rps13a) at S237 (***Figure 2L***) and Rps3a at S238 (***Figure 2M***) were entirely dephosphorylated by both Ism1 and insulin. Lastly, ribosomal protein S6 kinase alpha-5 (Rps6ka5), a known mTOR substrate (Chauvin et al., 2014) was phosphorylated by both Ism1 and insulin at S375 (***Figure 2N***). These data conclude that Ism1 induces phosphorylation of a specific set of proteins involved in mTOR signaling and proteostasis, while other phosphosites are unchanged (***Figure 2O***).

**Figure 2.**
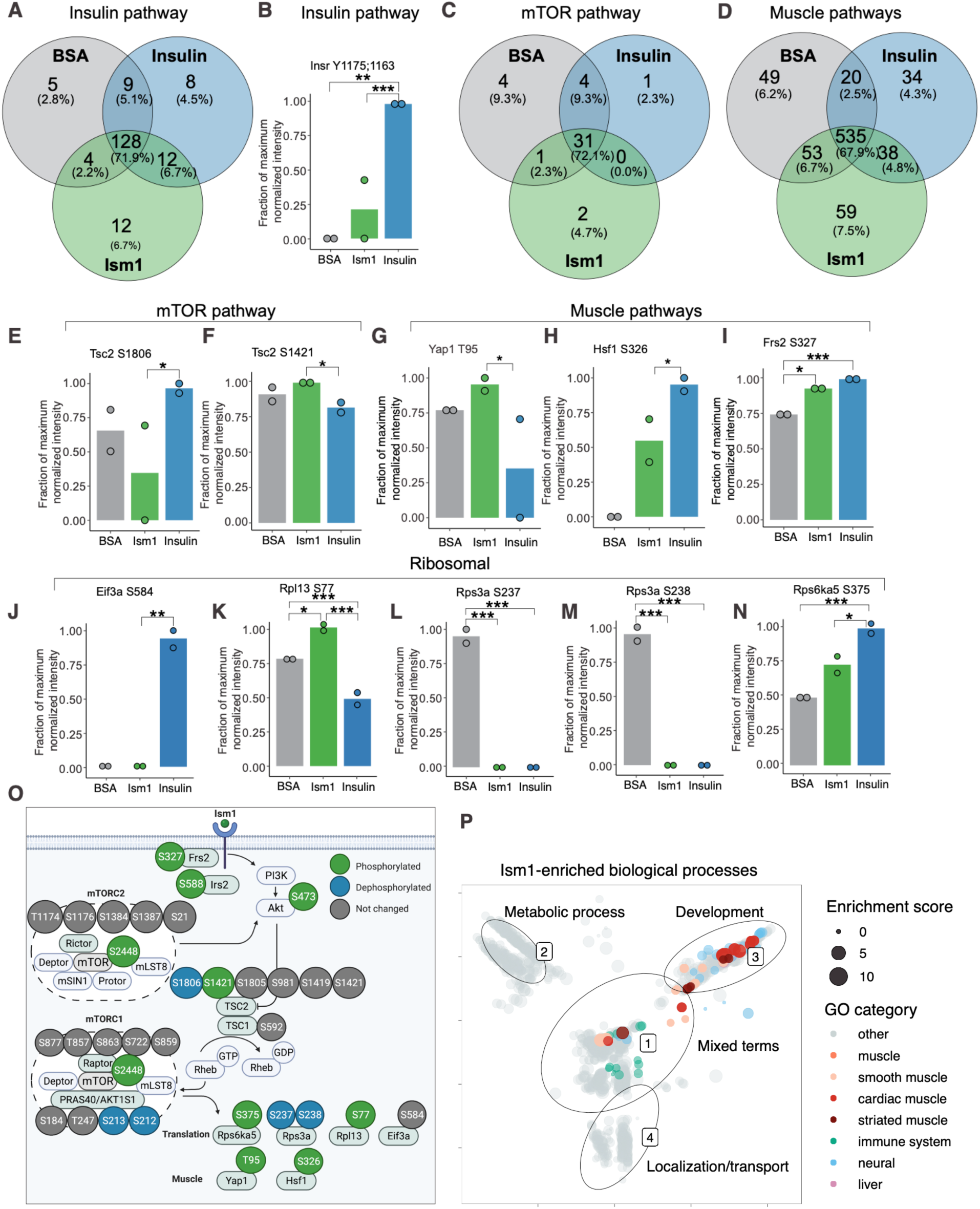
Phospho-specific mapping identifies an Ism1-induced signature consistent with protein translation and muscle function. (A) Venn diagram of shared and unique phosphosites between treatments for the GO pathways Insulin. (B) Abundance of InsR Y1175/1163 in cells treated with BSA, Ism1, or insulin (*n* = 2). Individual comparisons between conditions across phosphopeptides was performed using empirical Bayes statistics followed by adjustment for multiple testing using false discovery rate, *p < 0.05, **p < 0.01, ***p < 0.001. The minimum normalized intensity across the dataset was subtracted from each normalized data point, and phosphorylation was calculated as a fraction of the maximum value of all samples for each phosphopeptide. Bars show mean ± SEM. C) Venn diagram of shared and unique phosphosites between treatments for the GO pathways Insulin mTOR. D) Venn diagram of shared and unique phosphosites between treatments for the GO pathways and Muscle. (E-N) Abundance of proteins with indicated phosphosite in cells treated with BSA, Ism1, or insulin (*n* = 2). Individual comparisons between conditions across phosphopeptides was performed using empirical Bayes statistics followed by adjustment for multiple testing using false discovery rate, *p < 0.05, **p < 0.01, ***p < 0.001. The minimum normalized intensity across the dataset was subtracted from each normalized data point, and phosphorylation was calculated as a fraction of the maximum value of all samples for each phosphopeptide. Bars show mean ± SEM. O) Ism1 signaling network in 3T3-F442A cells. Ism1 ligand stimulation triggers activation of the PI3K/AKT pathway and the mTORC1 pathway which leads to changes in phosphorylation status of multiple proteins involved in protein translation and muscle function. P) Pathway analysis of enriched GO pathways in the Ism1 group versus BSA. Clusters are dominated by 1) mixed terms, 2) metabolic process, 3) development and 4) localization/transport. Plotted GO terms all have P values <0.01 calculated using the classic Kolmogorov-Smirnov test. See also Figure 2—figure supplement 1, and Figure 2—source data 1 and 2.

To globally discern possible pathways activated by Ism1 and possible grouping of functional effects, we conducted GO enrichment analysis using biological processes coupled with visualization by semantic similarity. We found associations previously linked to Ism1, including metabolic processes such as glucose and lipid metabolism (Jiang et al., 2021; Ruiz-Ojeda et al., 2022), nervous system development (Osoŕio et al., 2014; Pera et al., 2002), and the immune system (Lam et al., 2022; Li et al., 2021; Valle-Rios et al., 2014; Wu et al., 2021) (***Figure 2P***). Intriguingly, also here, we identify several pathways associated with muscle, skeletal muscle, and cardiac muscle in the Ism1 treatment group (***Figure 2P***). These results show a broad regulation of signatures indicating a role for Ism1 in muscle function.

### Ism1 induces mTOR-dependent protein synthesis in muscle cells

Given the Ism1-induced muscle-signaling signature in 3T3-F442A cells and that the PI3K-Akt pathway is known to promote anabolic programs in muscle cells (Edinger and Thompson, 2002), muscle cell differentiation (Wilson and Rotwein, 2007), and skeletal muscle hypertrophy (Bodine et al., 2001; Jaiswal et al., 2019), we next asked whether Ism1 induces anabolic cellular signaling pathways in muscle cells. In differentiated C2C12 myotubes, Ism1 induces phosphorylation of Akt^S473^ and ribosomal S6^S235/S236^ starting at 5 min and remaining up to 4h (***Figure 3A***). The effect of Ism1 on Akt signaling is robust but lower than that of the skeletal muscle hypertrophy hormone insulin-like growth factor-1 (Igf1) (***Figure 3A-B***). Similarly, undifferentiated C2C12 myoblasts are also responsive to Ism1 in a dose-dependent manner (***Figure 3C***). Notably, Ism1 treatment induced a 2.5-fold increase in protein synthesis as determined by [^35^S]-methionine incorporation into proteins (***Figure 3D***). As expected, Igf1 treatment resulted in a 3-fold induction in protein synthesis (***Figure 3D***), and the combined Igf1 and Ism1 treatment did not induce protein synthesis further, suggesting that the maximal capacity of protein synthesis has been reached under these conditions. Previous data showed that mTORC1/2 inhibition with rapamycin inhibits Ism1-induced signaling in 3T3-F442A cells, demonstrating that intact mTOR activity is required for the signaling capacity of Ism1 (Jiang et al., 2021). Importantly, low-dose rapamycin also inhibits Ism1-induced protein synthesis, establishing that the functional effects are directly linked to the signaling cascade induced by Ism1 (***Figure 3E***). These data align with the inhibitory effects of rapamycin on muscle hypertrophy during anabolic conditions (Pallafacchina et al., 2002). In conclusion, Ism1 induces a signaling cascade that requires intact mTOR signaling to induce protein synthesis in muscle cells.

**Figure 3.**
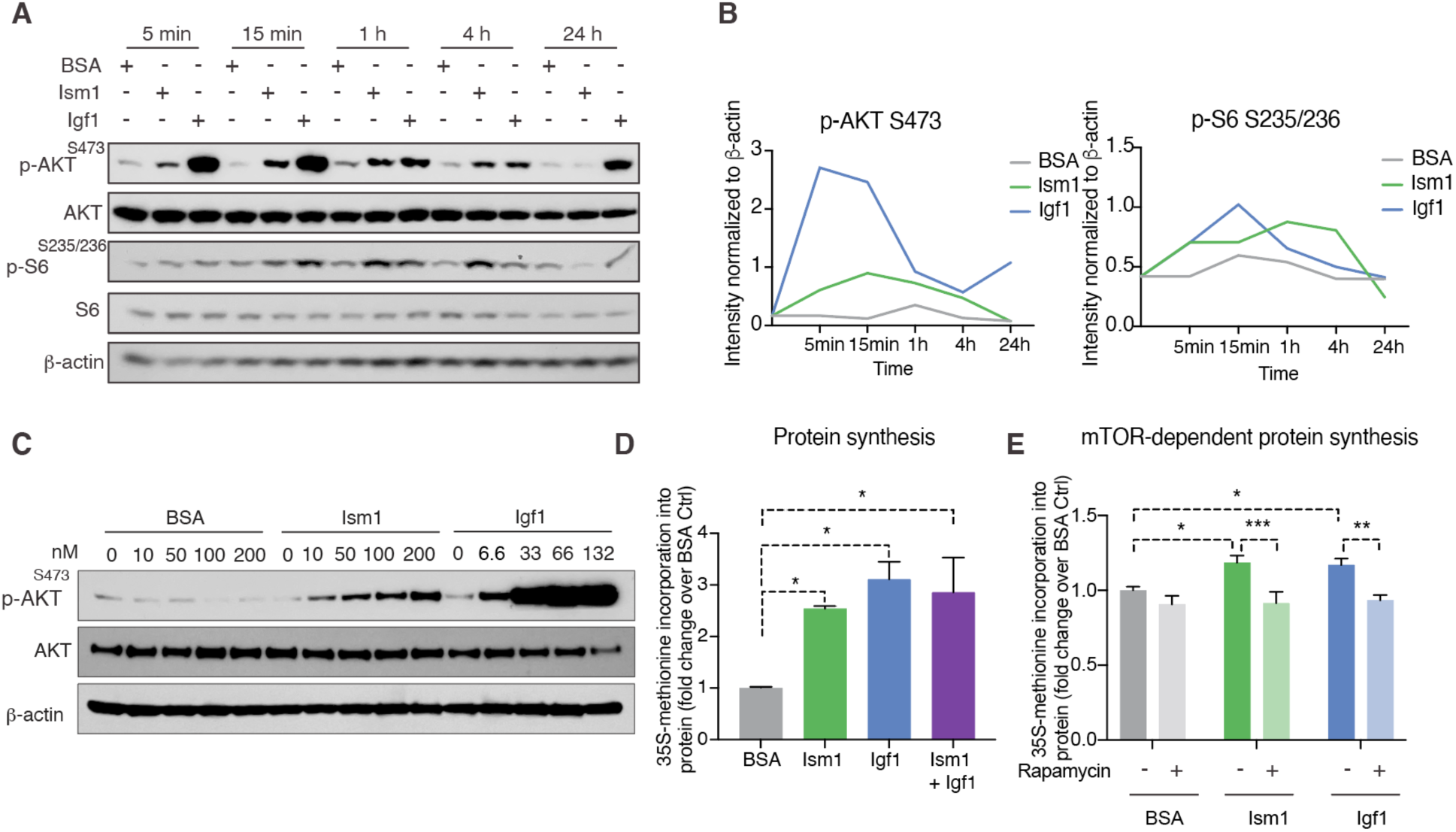
Ism1 induces mTOR-dependent protein synthesis in muscle cells. (A) Western blot analysis of p-AKT^S473^, total AKT, p-S6^S235/^236, total S6, and β-actin in C2C12 myotubes treated with 100 nM BSA, 100 nM Ism1, or 50 ng/ml Igf1. (B) Quantification of protein expression of p-AKT S473/ β-actin and p-S6 S235/236/β-actin. (C) Western blot analysis of p-AKT^S473^, total AKT and β-actin in C2C12 myoblasts treated with indicated concentrations of BSA, Ism1, or Igf1 for 5 minutes. (D) Levels of protein synthesis measured by [35S]-methionine incorporation in C2C12 cells after 48 hours of indicated treatments (*n* = 3, one-way ANOVA, *p < 0.05, **p < 0.01, ***p < 0.001). (E) Levels of protein synthesis measured by [35S]-methionine incorporation in C2C12 cells with indicated treatments for 2 hours in the presence or absence of 100 nM mTOR inhibitor rapamycin (*n* = 3, two-way ANOVA, *p < 0.05, **p < 0.01, ***p < 0.001). Bar graphs show mean ± SEM. See also Figure 3—source data 1 and 2.

### Ism1 ablation results in reduced skeletal muscle fiber size and muscle strength

Skeletal muscle atrophy, a reduction in muscle mass, occurs when the protein degradation rate exceeds protein synthesis (Cohen et al., 2015; Jaiswal et al., 2019; Sandri et al., 2004). We previously showed that Ism1 controls glucose uptake into adipose tissue and skeletal muscle in mice (Jiang et al., 2021), but the role of Ism1 in skeletal muscle function beyond glucose regulation has not been studied. Ism1 is broadly expressed, including highly in adipose tissue and blood (Jiang et al., 2021). By analyzing single-cell RNA sequencing data from murine skeletal muscle (Baht et al., 2020), we find that Ism1 is not expressed in muscle precursors or mature muscle cells (***Figure 4 - figure supplement 1***). Given the observed signaling action of Ism1 on muscle cells, this indicated a non-cell-autonomous effect of Ism1 on muscle cells in a physiological setting. Therefore, we next sought to determine whether whole-body ablation of Ism1 in mice, in which blood levels of Ism1 are completely ablated (Jiang et al., 2021), resulted in changes to skeletal muscle mass. To evaluate muscle atrophy, eight-week-old WT and Ism1-KO mice were either fasted for 12 h or fasted followed by a 12 h re-feeding period (fed state) (***Figure 4A***). As expected, at dissection, all fasted mice had a reduction in body weight compared with fed mice (***Figure 4B***). Blood glucose levels were 150 mg/dl in the fed state, and < 80 mg/dl in the fasted state (***Figure 4C***), but no differences were seen between genotypes. Under the same conditions, quadriceps (***Figure 4D***), gastrocnemius (***Figure 4 - figure supplement 2***), as well as tibialis (***Figure 4 - figure supplement 2***) muscles were harvested from mice under fed or fasted states and analyzed for muscle fiber size. Muscle tissue morphology was similar between the genotypes, but the muscle fiber size was notably smaller in the Ism1-KO quadriceps muscles (***Figure 4D***). Remarkably, fiber size area quantifications showed that the Ism1-KO quadriceps muscles demonstrate a robust shift in distribution to smaller muscle fiber size areas under both fed and fasted conditions (***Figure 4E***). On average, loss of Ism1 significantly reduces the cross-sectional area by 40 % in the fed state and > 20 % in the fasted state (***Figure 4F***). Given that the Ism1 phenotype was more pronounced under the fed state, we next evaluated the effect of Ism1 ablation on muscle function. While the heart weights (***Figure 4 - figure supplement 2***), muscle weights (***Figure 4 - figure supplement 2***), or total body weights (***Figure 4B***) of Ism1-KO mice were not significantly different from WT, the reduction in muscle fiber size is associated with impaired muscle force grip strength (***Figure 4G***). In conclusion, these results show that Ism1 ablation leads to smaller skeletal muscle fiber size and loss of muscle strength.

### Ism1 ablation does not impair movement or mitochondrial biogenesis, or normal muscle development

Since Ism1 ablation causes mild muscle atrophy, we next investigated whether loss of Ism1 is associated with a reduction in movement or muscle mitochondrial complex dependent mitochondrial bioenergetics, as has been demonstrated for IR and IGF-1R (Bhardwaj et al., 2021). Ambulatory activity (***Figure 5A***), respiratory exchange ratio (RER) (***Figure 5B***), energy expenditure as measured by VO_2_ consumption (***Figure 5C***), or food intake (***Figure 5D***) are indistinguishable between WT and Ism1-KO mice. Furthermore, protein levels of the mitochondrial complexes under fed and fasted conditions show no significant differences in any of the mitochondrial OXPHOS complexes I, II, III, IV, or V (***Figure 5E-F***). These data are consistent with the notion that the expression of the canonical regulator of mitochondrial biogenesis, *Ppargc1a* (*Pgc1α*) (Lin et al., 2002; Wu et al., 1999) is unchanged in quadriceps muscle tissues from the Ism1-KO mice (***Figure 5G***). General muscle markers, including the myosin heavy chain proteins *Myh1*, *Myh2*, *Myh4*, and *Myh7* were not altered under fed conditions, and only *Myh4* was reduced in the Ism1-KO mice under fasted conditions, suggesting that muscle differentiation is largely unaffected by Ism1 ablation and that the major role is in regulating hypertrophy in terminally differentiated muscle cells (***Figure 5H***). Taken together, these data suggest that Ism1 regulates muscle fiber size and muscle strength without affecting the expression of mitochondrial complexes or whole-body energy expenditure.

**Figure 4.**
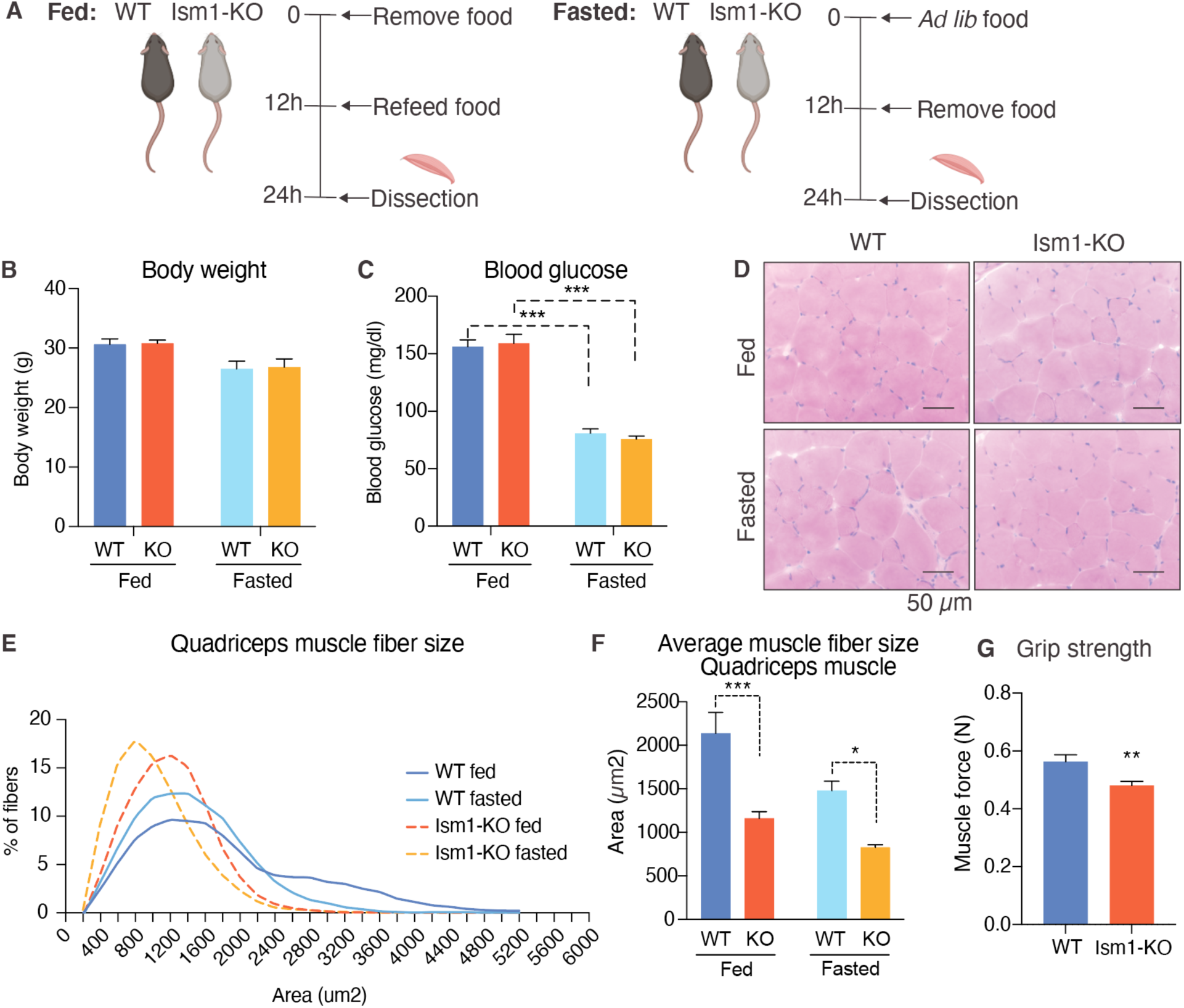
Ism1 ablation results in reduced skeletal muscle fiber size and muscle strength. (A) Schematic description of the fasting and feeding protocol. (B) Body weights of WT and Ism1-KO mice in the fed or fasted groups (WT fed, *n* = 3; Ism1-KO fed, *n* = 3; WT fasted, *n* = 3; Ism1-KO fasted, *n* = 3, two-way ANOVA). (C) Blood glucose level in fed and fasted mice before dissection (WT fed, *n* = 3; Ism1-KO fed, *n* = 3; WT fasted, *n* = 3; Ism1-KO fasted, *n* = 3, two-way ANOVA, *p < 0.05, **p < 0.01, ***p < 0.001). (D) H&E staining of mouse quadriceps muscles from WT and Ism1-KO mice in the fed or fasted groups (scale bars: 50 μm). Images are representative examples of three mice showing similar results. (E) Fiber size distribution of mouse quad muscles (WT fed, *n* = 3; Ism1-KO fed, *n*= 3; WT fasted, *n* = 3; Ism1-KO fasted, *n* = 3). Mean numbers of myotubes within the indicated area range is shown. (F) Quantification of average muscle fiber area (WT fed, *n* = 3; Ism1-KO fed, *n* = 3; WT fasted, *n* = 3; Ism1-KO fasted, *n* = 3, two-way ANOVA, *p < 0.05, **p < 0.01, ***p < 0.001) performed in a blinded fashion by two independent investigators. This experiment was repeated using two independent cohorts of mice. (G) Grip strength measured by muscle force (*N*) in WT and Ism1-KO mice (WT, *n* = 15; Ism1-KO, *n* = 15). P values are calculated by two-tailed Student’s t-test, *p < 0.05, **p < 0.01, ***p < 0.001. Bar graphs show mean ± SEM. See also Figure 4—figure supplement 1 and 2, and Figure 4—source data 1.

**Figure 5.**
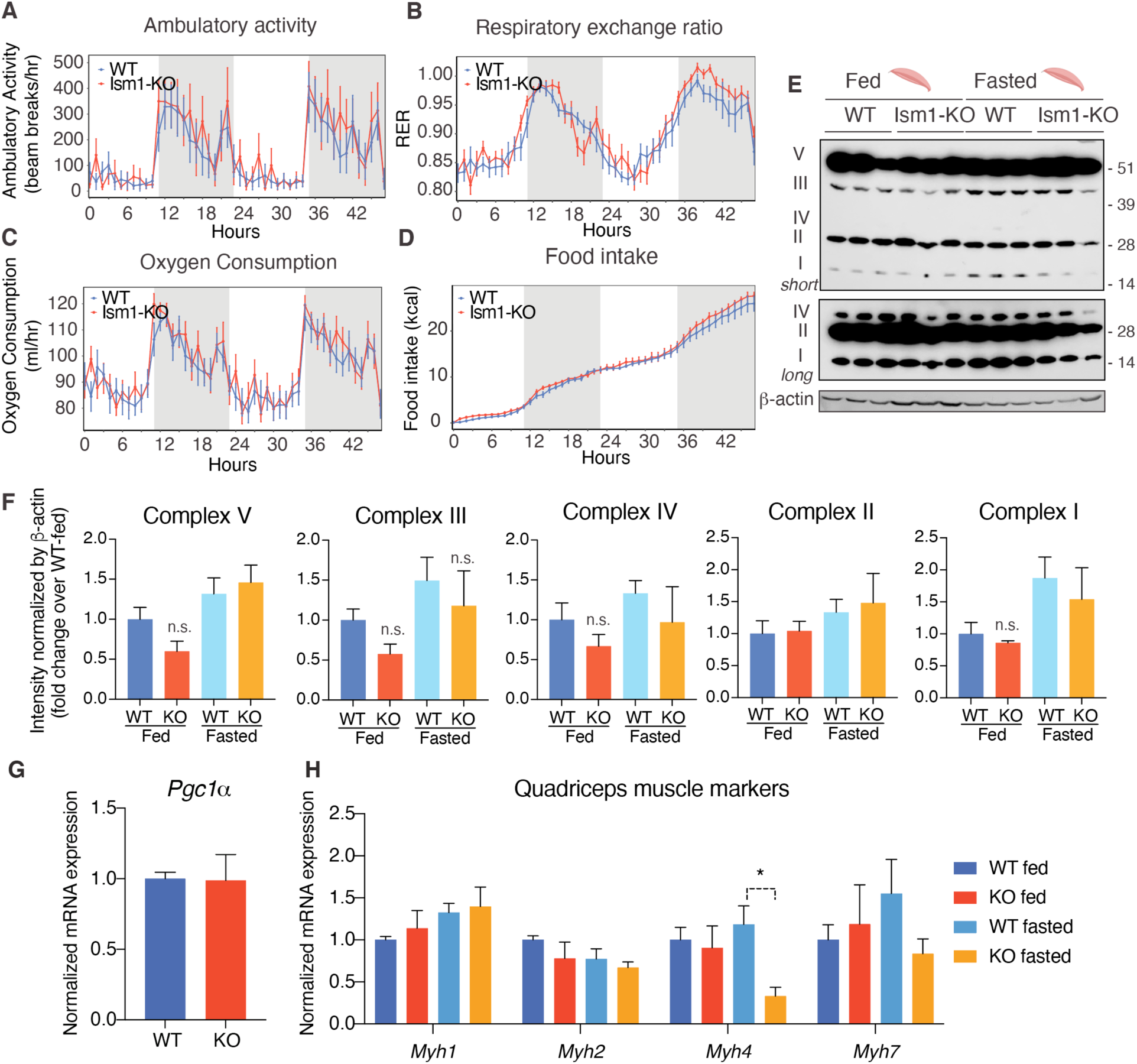
Ism1 ablation does not impair movement, mitochondrial biogenesis, or normal muscle development. (A) Ambulatory activity (WT, *n* = 4; Ism1-KO, *n* = 4, ANOVA, *p < 0.05), (B) Respiratory exchange ratio (RER) (WT, *n* = 4; Ism1-KO, *n*= 4, ANOVA, *p < 0.05), (C) Oxygen consumption (WT, *n* = 4; Ism1-KO, *n*= 4, ANCOVA, *p < 0.05), and (D) Food intake (WT, *n* = 4; Ism1-KO, *n*= 4, ANCOVA, *p < 0.05) in WT and Ism1-KO mice. (E) Levels of mitochondrial oxidative phosphorylation proteins in ETC complexes (OXPHOS) from quadriceps muscles (WT fed, *n* = 3; Ism1-KO fed, *n*= 3; WT fasted, *n* = 3; Ism1-KO fasted, *n* = 3) analyzed by Western blot. (F) Quantification of OXPHOS complexes (two-way ANOVA, *p < 0.05, **p < 0.01, ***p < 0.001). (G) Relative gene expression analysis of *Pgc1*α in quadriceps muscle from WT (*n* = 6) or Ism1-KO (*n* = 5) mice (two-tailed Student’s t-test, *p < 0.05, **p < 0.01, ***p < 0.001). (H) Relative gene expression analysis of *Myh1, Myh2, Myh4*, and *Myh7* in quadriceps muscle from WT fed (*n* = 6) or Ism1-KO fed (*n* = 5) vs. WT fasted (*n* = 5) or Ism1-KO fasted (*n* = 5) mice (two-way ANOVA, *p < 0.05, **p < 0.01, ***p < 0.001). Bar graphs show mean ± SEM. See also Figure 5—source data 1 and 2.

### Ism1-KO mice have defective skeletal muscle Akt and mTOR signaling and protein synthesis

To understand the underlying mechanism by which Ism1-KO mice develop smaller muscle fiber size, we next asked whether Ism1 directly controls protein content in mice. Quadriceps muscles from eight-week-old mice were harvested and analyzed for total protein content, demonstrating a significant reduction in total protein content in the Ism1-KO mice compared with WT mice (***Figure 6A***). Because muscle proteostasis is balanced by protein synthesis and degradation (Glass, 2011; Jackman and Kandarian, 2004), we next tested the hypothesis that Ism1 is required for efficient protein synthesis in mice by measuring *in vivo* [^35^S]-methionine incorporation into proteins isolated from quadriceps muscles (***Figure 6B***). WT and Ism1-KO mice were I.P. administered with [^35^S]- methionine for 2 hours, followed by protein precipitation, which demonstrates that loss of Ism1 results in a significant reduction in muscle protein synthesis in mice (***Figure 6C***). Furthermore, under either fed or fasted conditions, muscles isolated from Ism1-KO mice have significantly increased levels of atrophy genes FoxO1 (***Figure 6D***), the FoxO target genes *Cdkn1b*, *Eif4ebp1*, and *Ctsl* (***Figure 6E***), and ubiquitin ligase *Fbxo30* compared with WT mice (***Figure 6F***). These data suggest that Ism1 controls protein synthesis, and its loss is associated with elevated protein degradation gene expression. Given that Ism1 activates the Akt pathway in myocytes *in vitro*, and that loss of muscle-specific Akt1 and Akt2 leads to muscle atrophy (Jaiswal et al., 2019), we next hypothesized that the mechanism behind the smaller muscle fiber size in the Ism1-KO mice is due to reduced Akt pathway activity. Indeed, we found that phosphorylation of Akt^S473^, mTOR^2448^, and ribosomal S6^S235/S236^ are markedly decreased in Ism1-KO muscle compared with WT mice under both fed and fast conditions, providing an explanation for the lower protein synthesis rate (***Figure 6G***). Taken together, these results demonstrate that Ism1 is a circulating anabolic regulator of protein synthesis and muscle function by maintaining high Akt activity and protein synthesis in skeletal muscle. In conclusion, Ism1 is a circulating anabolic regulator of protein synthesis and muscle function.

**Figure 6.**
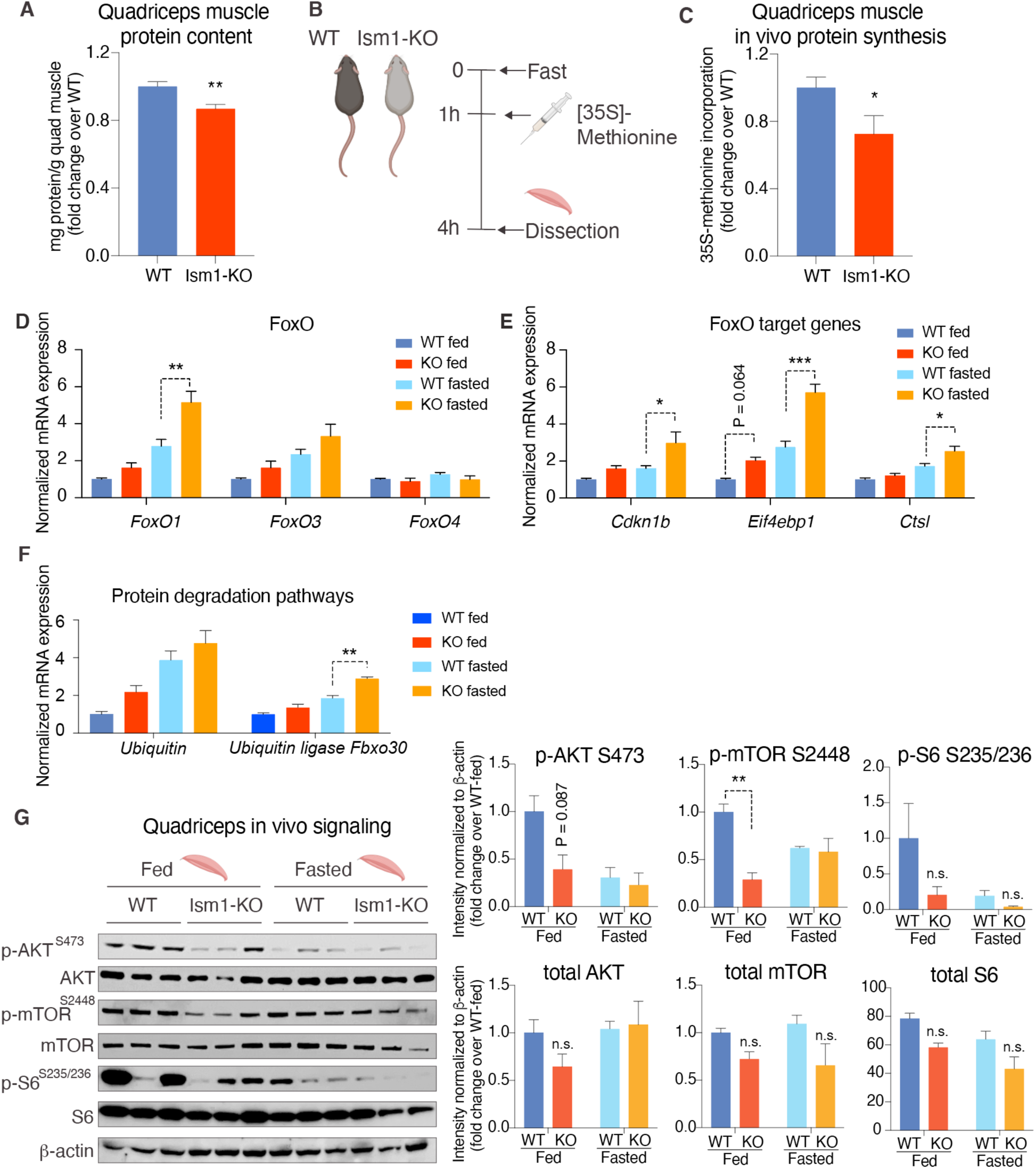
Ism1-KO mice have defective skeletal muscle protein synthesis and AKT-mTOR signaling. (A) Total protein content measured in WT and Ism1-KO quadriceps muscle expressed as mg protein/wet tissue weight in grams. (WT, *n* = 12; Ism1-KO, *n* = 9, two-tailed student’s t-test *p < 0.05, **p < 0.01, ***p < 0.001). (B) Schematic description of the *in vivo* [35S]-methionine incorporation protocol. (C) *In vivo* protein synthesis measured by [35S]-methionine incorporation in WT and Ism1-KO mice (WT, *n* = 12; Ism1-KO, *n* = 9, two-tailed Student’s t test, *p < 0.05, **p < 0.01, ***p < 0.001). Relative gene expression analysis of (D) *FoxO* and (E) *FoxO* target genes *Cdkn1b, Eif4ebp1,* and *Ctsl* in quadriceps muscle from WT fed (*n* = 6) or Ism1-KO fed (*n* = 5) vs. WT fasted (*n* = 5) or Ism1-KO fasted (*n* = 5) mice (two-way ANOVA, *p < 0.05, **p < 0.01, ***p < 0.001). (F) Relative gene expression analysis of *Ubiquitin* and the ubiquitin ligase *Fbxo30* in quadriceps muscle from WT fed (*n* = 6) or Ism1-KO fed (*n* = 5) vs. WT fasted (*n* = 5) or Ism1-KO fasted (*n* = 5) mice. P values are calculated by two-way ANOVA, *p < 0.05, **p < 0.01, ***p < 0.001. Bar graphs show mean ± SEM. (G) Western blot analysis and quantification of the *in vivo* levels of pAKT^S473^, total AKT, p-mTOR^S2448^, total mTOR, pS6^S235/236^, total S6, and β-actin in quadriceps muscles of WT and Ism1-KO mice under fed and fasted conditions (WT fed, *n* = 3; Ism1-KO fed, *n* = 3, WT fasted, *n* = 3; Ism1-KO fasted, *n* = 3, two-way ANOVA, *p < 0.05, **p < 0.01, ***p < 0.001). Bar graphs show mean ± SEM. See also Figure 6—source data 1 and 2.

## Discussion

Akt has an established role in enhancing muscle hypertrophy and function (Bodine et al., 2001; Glass, 2011; Jaiswal et al., 2022, 2019; Lai et al., 2004; Mammucari et al., 2007; Wilson and Rotwein, 2007). However, there is still a need to identify other hormonal and physiological insulin/IGF-1 independent activators of Akt to avoid associated side effects such as hypoglycemia when used therapeutically. We previously identified Ism1 as a secreted protein that activates Akt in multiple cell types, including human skeletal muscle cells. Using radiolabeled glucose, we also showed that Ism1 increases glucose transport into both adipose tissue and skeletal muscle (Jiang et al., 2021). Several lines of evidence from the current study suggest that Ism1 also has an important anabolic role in promoting skeletal muscle growth. Most importantly, Ism1 acts directly on muscle cells to induce protein synthesis. Conversely, Ism1 ablation leads to lower skeletal muscle fiber size and atrophy associated with lower Akt signaling and protein synthesis. Ism1-KO mice have elevated levels of FoxO1 target genes consistent with increased protein degradation. Therefore, Ism1 appears to be an important protein regulating metabolism during fed and fasted conditions.

The signaling mechanism behind Ism1-induced muscle hypertrophy is not simply an activation of the identical insulin/IGF-1-induced PI3K-Akt pathway. Across multiple cell types, Ism1 and insulin share the pAkt and pS6 network but are seemingly segregated by the more robust activation of pAkt^S473^ seen with insulin, whereas Ism1 induces only a subset of shared Akt-induced insulin targets. Interestingly, while Ism1 does not directly phosphorylate the IR/IGFRs, Ism1 does induce phosphorylation of the insulin receptor substrate proteins (Irs), a feature that is shared with other endocrine hormones. This phosphospecific regulation by distinct hormones can result in a diverse range of functional outcomes (Yenush and White, 1997). For example, Irs1 phosphorylation at S302, S307, S522 and S636/639 have been linked to insulin resistance (Um et al., 2004), but not all hormones that phosphorylate those Irs sites induce insulin resistance, including FGF21 (Minard et al., 2016). These overlapping but distinct pathways may account for the Ism1’s cell-type specific functional outcomes and downstream transcriptional effects unique to Ism1. Additionally, the phosphoproteomic mapping shows a distinct muscle signature specifically induced by Ism1 and not by insulin. Among Ism1 downstream targets, we find phosphorylation of proteins related to the mTOR pathway, ribosomal and muscle function, targets not previously identified downstream of Ism1.

In this work, we observe a 60 % decrease in pAkt^S473^ levels in the muscles of Ism1-KO mice, which leads to a ∼10 % reduction in muscle protein content. Our data is consistent with previous reports using muscle-specific Akt-KO mice that demonstrate a 40% reduction in protein synthesis upon complete ablation of Akt (Jaiswal et al., 2022). Furthermore, reduced Akt is associated with skeletal muscle insulin resistance, in line with our previous observation that Ism1-KO mice fed a high-fat diet are more insulin resistant (Jiang et al., 2021). To what extent the insulin resistance in the Ism1-KO mice contributes to muscle function in obesity or during aging is an intriguing question that remains to be answered in future work. Moreover, increasing protein synthesis while suppressing protein degradation is important in a physiological setting, based on the findings that only a combination of FoxO1 inhibition and mTORC1 activation can restore Akt-mediated muscle loss (Jaiswal et al., 2022). This suggests that hormonal regulators of Akt activity could have important biological functions because of Akt’s dual role in regulating mTORC1 and the FoxO genes. Here we used genetic and pharmacologic approaches to determine the role of Ism1 in regulating protein synthesis, but it remains to be explored whether pharmacological Ism1 administration prevents protein degradation in the skeletal muscle. Additionally, upon muscle hypertrophy, activated muscle precursors or satellite cells provide an additional mechanism for muscle expansion (Hawke and Garry, 2001). A limitation of this study is that we did not investigate whether Ism1 affects muscle regeneration. As IGF-1 induces the growth and differentiation of satellite cells (Musarò et al., 1999), and HGF induces activation of cell cycle G_Alert_ (Rodgers et al., 2017), it will be interesting to determine if Ism1 also activates skeletal muscle stem cells. Identifying all physiological mechanisms that distinguish Ism1 from IGF-1 will aid in understanding whether other yet unidentified pathways distinct from Akt and FoxO regulate proteostasis.

The physiological function of Ism1 in regulating muscle growth is expected given that Ism1 stimulates Akt, but it was somewhat unexpected that the lower muscle protein content and fiber size did not lead to a significant loss of muscle mass. It remains to be determined what replaces protein mass in the muscle of Ism1-KO mice, including water, glycogen stores, or lipids, which could be regulated by Ism1. Notably, Ism1 itself is not expressed in muscle cells but only in other cells, including adipocytes and immune cells. Circulating levels of human ISM1 using immunoreactive ELISAs have been reported to be in the range of 1-50 ng/ml (Jiang et al., 2021; Ruiz-Ojeda et al., 2022), consistent with mass spectrometry peptide analysis which estimates a circulating ISM1 concentration of 17 ng/ml (Uhlén et al., 2019). The necessary circulating and local Ism1 levels to maintain muscle fiber size or prevent fasting-induced muscle loss remain to be determined. It is plausible that ISM1 levels are altered in response to fasting and feeding, insulin resistance, or aging - conditions associated with temporary or chronic muscle loss (Laurens et al., 2021; Perry et al., 2016; Roubenoff and Castaneda, 2001). A limitation of the current study is that fasting, but not more severe atrophy conditions were studied. Long-term studies ought to explore if pharmacological administration of Ism1 systemically or locally into the muscle is sufficient to prevent muscle loss in more severe models of muscle loss. Nevertheless, as aging impairs skeletal muscle protein synthesis and leads to muscle weakness and atrophy, the current work has implications for multiple conditions, including diabetes- and age-induced muscle atrophy.

## Materials and Methods

### Animal studies

Animal experiments were performed per procedures approved by the Institutional Animal Care and Use Committee of the Stanford Animal Care and Use Committee (APLAC) protocol number #32982. C57BL/6J mice were purchased from the Jackson Laboratory (#000664). The Ism1-KO mice (C57BL/6J-Ism1^em1Kajs^/J strain #036776, JAX) were generated using the Ism1 floxed allele as described previously (Jiang et al., 2021). Unless otherwise stated, mice were housed in a temperature-controlled (20-22°C) room on a 12-hour light/dark cycle. All experiments were performed with sex- and age-matched, littermate male mice housed in groups of five unless stated otherwise.

### Sample preparation for phosphoproteomics analysis

The phosphoproteomics analysis was performed using 3T3-F442A cells. Cells were cultured in DMEM/F12 medium with 10 % FBS/1 % pen/strep until 80-100 % confluent. Following a 16 h starvation in serum-free DMEM/F12 medium, cells were treated with PBS, 100 nM Ism1 or 100 nM insulin for 5 minutes (*N* = 6 per treatment group, 10M cells/treatment). Following treatment, the medium was aspirated, and cells were washed three times with ice-cold PBS while kept on ice. 1 ml ice-cold PBS supplemented with complete mini protease inhibitors (#4693124001, Sigma-Aldrich, St. Louis, MO, USA) and phosSTOP phosphatase inhibitors (#4906845001, Sigma-Aldrich, St. Louis, MO, USA) were added to the cells that were immediately scraped down, pelleted at 14,000 *x* g for 10 min followed by snap freezing. Cell pellets were kept at -80°C until analysis. Aliquots of the samples were used to verify AKT^S473^ signaling induction by Ism1 and insulin by western blot. Six biological replicates per treatment group were pooled into two replicates per treatment group for the phosphoproteomics analysis. The phosphoproteomics analysis was performed at Northwestern University Proteomics core as described previously (Dephoure et al., 2013). Briefly, protein extracts were alkylated with iodoacetamide and digested with trypsin at a ratio of 1:50 (w/w) trypsin to protein. Tryptic phosphopeptides were enriched by TiO_2_-immobilized metal affinity chromatography, washed with 70 % (v/v) EtOH and equilibrated in 1 % Nh4OH and three times in 1M glycolic acid in 80 % (v/v) acetonitrile, 5% (v/v) trifluoroacetic acid. Peptides were eluted and dried in a SpeedVac concentrator. Peptides were resuspended in 3 % (v/v) acetonitrile, 0.1 % trifluoroacetic acid and twice in 0.1 % trifluoroacetic acid. Dried peptides were dissolved in LC-MS/MS solvent (3 % acetonitrile and 0.1 % fluoroacetic acid prior to LC-MS/MS analysis using a dual-pressure linear ion trap (Velos Pro) with a high-field Orbitrap mass analyzer as described previously (Yue et al., 2015; Zhou et al., 2008).

### Phosphoproteomics analysis

Data acquired on the Orbitrap were searched against a Uniprot *Mus Musculus* protein database using TDPortal as previously described (Fornelli et al., 2017). Annotations were extracted from UniProtKB, Gene Ontology (GO), and the Kyoto Encyclopedia of Genes and Genomes (KEGG). Bioinformatics analyses including hierarchical clustering, pathway analysis and annotation enrichment were performed with R v4.1.1 using the DEP package (Zhang et al., 2018) for normalization and testing for significance. A varying number of permutations of imputations and testing for significance were applied until the insulin receptor was consistently found to be phosphorylated by insulin versus BSA. Pathway enrichment was analyzed using the TopGO package using the classic Kolmogorov-Smirnov test. We applied semantic similarity analysis between GO terms followed by principal coordinates analysis to cluster and visualize enrichments. Unless otherwise stated, individual comparisons between conditions across phosphopeptides was performed using empirical Bayes statistics followed by adjustment for multiple testing with a p-value of * < 0.05, **p < 0.01, ***p < 0.001 to be considered significant. The minimum normalized intensity across the dataset was subtracted from each normalized data point, and phosphorylation was calculated as a fraction of the maximum value of all samples for each phosphopeptide. Plotted GO terms have P values <0.01 calculated using the classic Kolmogorov-Smirnov test. The distribution diagram of shared and unique phosphosites was obtained by retrieving the selected GO pathways for peptides detected in at least 1 sample between treatments. Specific details on the significance of each test are described in the figure legends.

### Immunohistochemistry and muscle fiber size quantification

Tissues were snap-frozen and cross-sectioned at 10 μm. For hematoxylin and eosin (H&E) staining, slides were stained with hematoxylin for 3 min, washed with water and 95% ethanol, and stained with eosin for 30 min. Sections were then washed with ethanol and xylene, and mounted with mounting medium. The tissue slides were observed with a Nikon 80i upright light microscope using a 20 x objective lens. Digital images were captured with a Nikon Digital Sight DS-Fi1 color camera and NIS-Elements acquisition software. Muscle fiber sizes for the pectoralis, quadriceps, soleus, gastrocnemius, and tibialis muscle tissues were determined by measuring cross-sectional area (μm2) using the Image J (version 1.53e) software. The muscle fibers were manually outlined to obtain their measurement data. Blind scoring by two independent investigators of the muscle tissues was done to unbiasedly collect data for all categories of mice. The quantification of the average muscle fiber area was performed on *N* = 3 independent muscle tissues from each genotype, with 2-4 photos taken from each muscle tissue and average values calculated for each photo. Statistically significant differences were determined using two-way ANOVA.

### Indirect calorimetry, food intake and physiological measurements

Oxygen consumption (VO_2_), respiratory exchange ratio, movement, and food intake in eight-to-twelve-week-old WT and Ism1-KO mice were measured using the environment-controlled home-cage CLAMS system (Columbus Instruments, Columbus, OH, USA) at the Stanford Diabetes Research Center. Mice were maintained on a chow diet and housed at 20-22°C in the cages for 24 h prior to the recording. Energy expenditure calculations were not normalized for body weight. RER and locomotion were analyzed using ANOVA. All other calorimetry measurements in mice were analyzed using ANCOVA using the CalR version 1.3 without the remove outliers feature (Mina et al., 2018). The muscle strength in the hindlimbs was measured with a grip meter. The mice were trained to grasp a horizontal bar while being pulled by their tail. The force (expressed in Newton) was recorded by a sensor.

### Single-cell RNA sequencing of skeletal muscle in mice

Single-cell RNA sequencing data was re-analyzed from a previously published dataset performed on single-cell suspension from murine tibialis anterior skeletal muscles (Baht et al., 2020). Briefly, tibialis anterior muscles from three injured mice were pooled as well as three uninjured mice to generate two samples used for scRNA-Seq. Three thousand single cells from each of the two samples were barcoded and cDNA generated using 10X Genomics Chromium Drop-seq platform and sequencing on Illumina 2500 platform. 10X Genomics Cell Ranger software was used to demultiplex and align reads. Seurat (V4.2) package was used to perform quality control, sample normalization and clustering for cell type identification. Downstream analyses included gene analysis of the genes of interests associated with cell clusters.

### Expression and purification of recombinant proteins

The Ism1 proteins were generated by transient transfection of mouse Ism1 with C-terminal Myc-6X-his tag DNA plasmids Addgene (#173046) into Expi293F cells. Recombinant proteins were produced in mammalian Expi293F cells using large-scale transient DNA transfection and purified using Cobalt columns and buffer exchanged to PBS. Protein purity and integrity were assessed with SDS page, Superdex200 size exclusion column and endotoxin assay. Every protein batch produced was tested for bioactivity by measuring the induction of pAKT^S473^ signaling in 3T3-F442A cells as described previously (Jiang et al., 2021). All proteins were aliquoted and stored at -80 °C and not used for more than three freeze-thaws.

### Culture and differentiation of C2C12 cells

C2C12 cells (#CRL-1772, ATCC) were cultured in DMEM with 10 % FBS. Cells were passaged every two days and were not allowed to reach more than 70 % confluency. For differentiation, media was replaced with DMEM with 2 % horse serum and cultured for four days to induce myotube formation within 4-6 days as described previously(Risson et al., 2009; Sandri et al., 2004).

### *In vivo* and *in vitro* protein synthesis

*In vivo* protein synthesis was measured by incorporation of [^35^S]-methionine into proteins isolated from skeletal muscle in mice. Briefly, mice fasted for 1h were I.P. injected with 2.5 μCi/gram [^35^S]-methionine diluted in saline. Two hours after injection, the quadriceps muscles were removed, weighed, and snap-frozen in liquid nitrogen. The tissues were homogenized using a hand-held homogenizer in RIPA buffer containing protease inhibitor cocktail (Roche) and centrifuged at 4 °C to remove cell debris. Protein concentration in the supernatant was determined by BCA assay (Thermo Fischer Scientific, Waltham, MA, USA), and total protein content was calculated by multiplying the protein concentration by the supernatant volume. Proteins were extracted using TCA precipitation and the radioactivity was counted on a scintillation counter. Protein synthesis in C2C12 cells was measured by incorporation of [^35^S]- methionine into proteins using a modified protocol developed for skeletal myotubes (Hong-Brown et al., 2007; Kazi and Lang, 2010; Méchin et al., 2007) as described previously (Schmidt et al., 2009). For [^35^S]-methionine incorporation, C2C12 cells were treated with BSA, recombinant Ism1 or Igf1 with the addition of 0.5 μCi [^35^S]-methionine (#NEG009L005MC, PerkinElmer, Waltham, MA, USA) for 48h. For [^35^S]-methionine incorporation in the presence or absence of inhibitors, C2C12 cells were treated with DMSO or 100 nM Rapamycin for 2h, followed by treatments with BSA, recombinant Ism1 or Igf1 for 1h. Subsequently, 0.5 μCi [^35^S]-methionine (#NEG009L005MC, PerkinElmer) was added for another 1h. To stop the incubation, cells were washed in ice-cold PBS three times. Proteins were extracted using TCA precipitation and the radioactivity was counted on a scintillation counter.

### Gene expression analysis

Total RNA from cultured cells or tissues was isolated using TRIzol (Thermo Fischer Scientific, Waltham, MA, USA) and Rneasy mini kits (QIAGEN, Hilden, Germany). RNA was reverse transcribed using the ABI high-capacity cDNA synthesis kit. For q-RT-pcr analysis, cDNA, primers and SYBR-green fluorescent dye (Bimake, Houston, TX, USA) were used. Relative mRNA expression was determined by normalization to Cyclophilin levels using the ΔΔCt method.

### Western blots and molecular analyses

For western blotting, homogenized tissues or whole-cell lysates, samples were lysed in RIPA buffer containing protease inhibitor cocktail (Roche, Basel, Switzerland) and phosphatase inhibitor cocktail (Roche, Basel, Switzerland), prepared in 4X LDS Sample Buffer (Invitrogen, Waltham, MA, USA) and separated by SDS-PAGE and transferred to Immobilon 0.45µm membranes (Millipore, Burlington, MA, USA). The antibodies used are as follows: Rabbit monoclonal anti-p-AKT1/2 (Ser473) (#4060), AKT1 (pan) (#4691 CST), rabbit monoclonal anti-p-mTOR (Ser2448) (D9C2), rabbit polyclonal anti-p-S6 Ribosomal Protein (Ser235/236) (Cat#2211 CST), ribosomal protein S6 (#2217 CST) from Cell Signaling. Mouse monoclonal anti-beta Actin AC-15 HRP (#AB49900) and OXPHOS rodent antibody (#ab110413) were from Abcam. Donkey anti-rabbit IgG (HRP) (#NA934) and sheep anti-mouse IgG (HRP) (#NA931) were from Cytiva (GE). Recombinant Igf1 (#791-MG-050) was from R&D systems. The mammalian expression plasmid for Ism1 with C-terminal myc-6xhis tag plasmid for recombinant Ism1 protein production was from Addgene (#173046).

### Statistical analyses

Values for *N* represent biological replicates for cultured cell experiments or individual animals for *in vivo* experiments. Group-housed mice within eight weeks of age were used for comparative studies. Mice were randomly assigned to treatment groups for *in vivo* studies. Significant differences between the two groups (* p < 0.05, ** p < 0.01, *** p < 0.001) were evaluated using a two-tailed, unpaired Student’s t-test as the sample groups displayed a normal distribution and comparable variance (Prism9 software; GraphPad). Two-way ANOVA with repeated measures was used for body weight and repeated measurements (* p < 0.05, ** p <0.01, *** p < 0.001). All data are presented as the standard error of mean (S.E.M.) or as described in the figure legend. Specific details for *N* values are noted in each figure legend.

## Acknowledgements

K.J.S. was supported by NIH grants DK125260, DK111916, the Stanford Diabetes Research Center P30DK116074, the Jacob Churg Foundation, the McCormick and Gabilan Award, the Weintz Family COVID-19 research fund, American Heart Association (AHA), the Stanford School of Medicine, and the Stanford Cardiovascular Institute (CVI). M.Z. was supported by the American Heart Association (AHA) postdoctoral fellowship (905674). L.V. was supported by Stanford School of Medicine Dean’s Postdoctoral Fellowship. E.B.M. was supported by Stanford School of Medicine Dean’s Postdoctoral Fellowship and the American Heart Association (AHA) postdoctoral fellowship (18POST34030448). D.E.L. was supported by NIH training grant T32HL007057. J.P.W. was supported by NIH/NIA grant K01AG056664 and R21AG065943 and Borden Scholar Award through Duke University. N.C. was supported by the American Heart Association (AHA) SURE award (882082). N.B.D.S. was supported by the Carlsberg Foundation Internationalization Fellowship. C.V.R. was supported by the Weintz family COVID-19 research fund, the Stanford Women’s Cancer Center and the Stanford Cancer Institute, The Jacob Churg award, and the Children’s Cancer Research Foundation. We thank the Stanford University Pathology Histology core facility, and the Pathology Department for microscopy equipment. We thank the Proteomics core at Northwestern University for assistance with the phosphoproteomic analysis, and Pratima Nallagatla at Stanford University for assistance with the data processing. This work was supported by the Stanford Diabetes Research Center (NIH grant #P30DK116074). Some illustrations were created with Biorender under a paid subscription.

## Additional information

### Author Contributions

Conceptualization, M.Z, and K.J.S.; Methodology, M.Z, N.B.D.S., K.J.S. Investigation, M.Z., N.B.D.S., L.V., L.U., E.B.M., D.E.L., J.P.W., N.C., Q.N., C.V.R., K.J.S., Writing - Original Draft, M.Z, N.B.D.S., K.J.S.; Writing - Review & Editing, M.Z, N.B.D.S., L.V., K.J.S.; Resources, C.V.R., K.J.S.; Supervision and funding acquisition, K.J.S.

### Additional files

**Source data 1**

Unedited raw western blot images in Figure 1, 3, 5 and 6.

**Figure 1—supplement 1.**
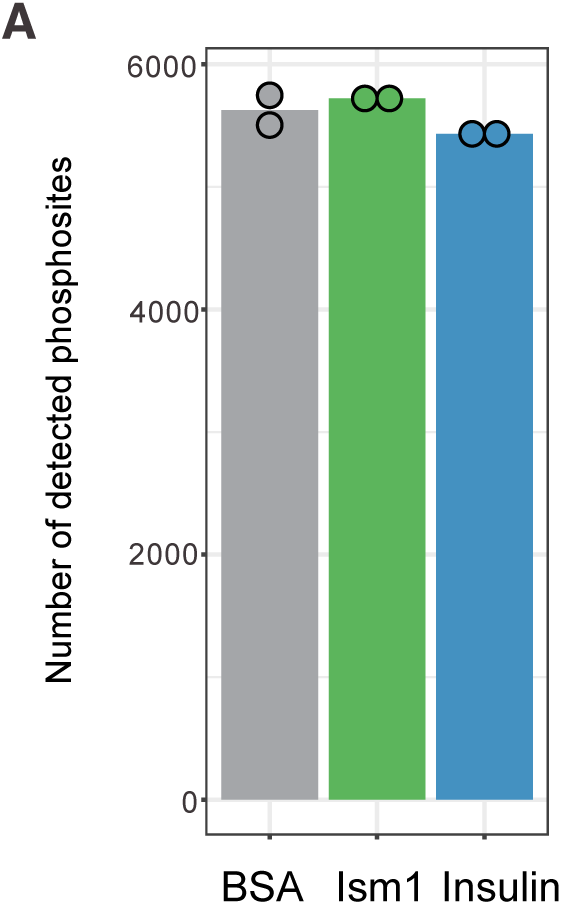
Quality controls for the phosphoproteomics analysis. (A) Total phosphosites identified in cells treated with 100 nM bovine serum albumin (BSA), 100 nM Ism1, or 100 nM insulin for 5 minutes.

**Figure 1—source data 1**

List of phosphosites significantly different between treatments for Figure 1E.

**Figure 1—source data 2**

List of shared and unique phosphosites between treatments for selected GO pathways for Figure 1F.

**Figure 1—source data 3**

Uncropped western blot images with relevant bands labelled.

**Figure 2—supplement 1.**
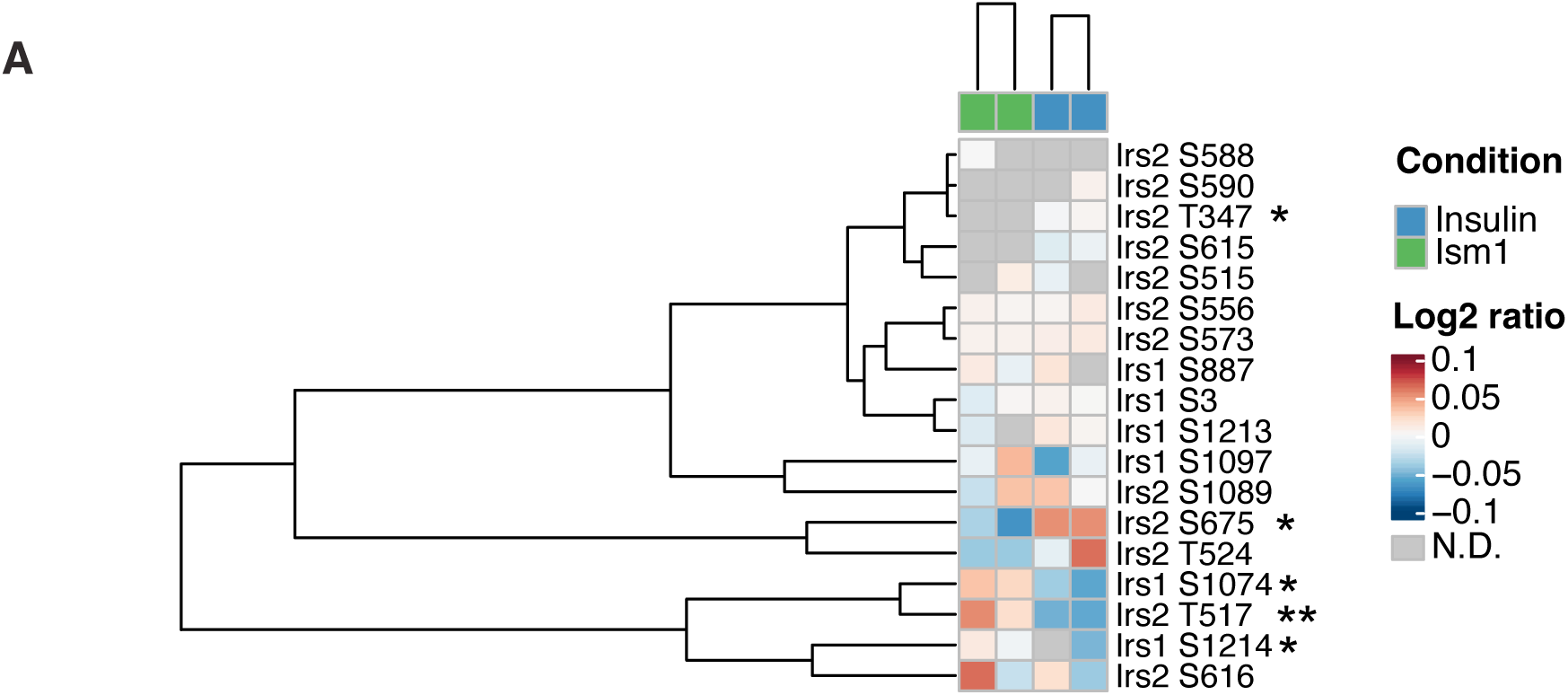
Insulin receptor substrate-1 and 2 (Irs1/2) phosphorylation status in response to Ism1 or insulin. (A) Heatmap of differentially phosphorylated Irs1 and Irs2 phosphosites by Ism1 relative to insulin (*n* = 2, individual comparisons between conditions across phosphopeptides using empirical Bayes statistics followed by adjustment for multiple testing using false discovery rate, *p < 0.05, **p < 0.01, ***p < 0.001).

**Figure 2—source data 1**

Enriched pathways for proteins with phosphosites significantly different between Ism1 and BSA clustered by semantic similarity.

**Figure 2—source data 2**

Raw data related to Figure 2B, and E-N.

**Figure 3—source data 1**

Raw data related to Figure 3B, D, and E.

**Figure 3—source data 2**

Uncropped western blot images with relevant bands labelled.

**Figure 4—supplement 1.**
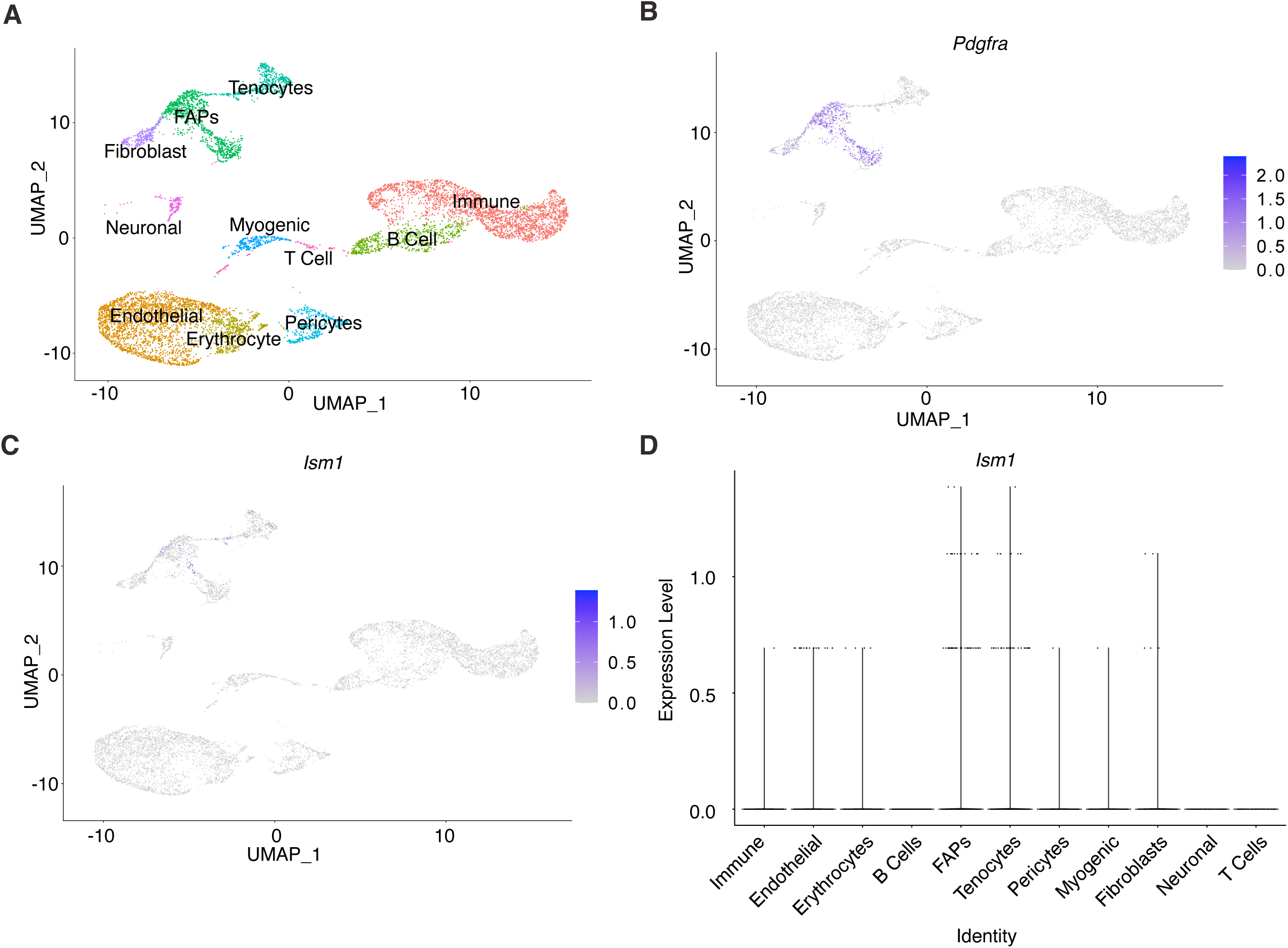
Ism1 acts non-cell autonomously on muscle cells. (A) UMAP plot of single-cell RNA sequencing of isolated cells from mouse skeletal muscle. (B) Expression of *Pdgfra* in fibro-adipose precursors within the muscle. (C) Low expression of *Ism1* within the muscle tissue. (D) Low expression of Ism1 in myocytes and myoblasts.

**Figure 4—supplement 2.**
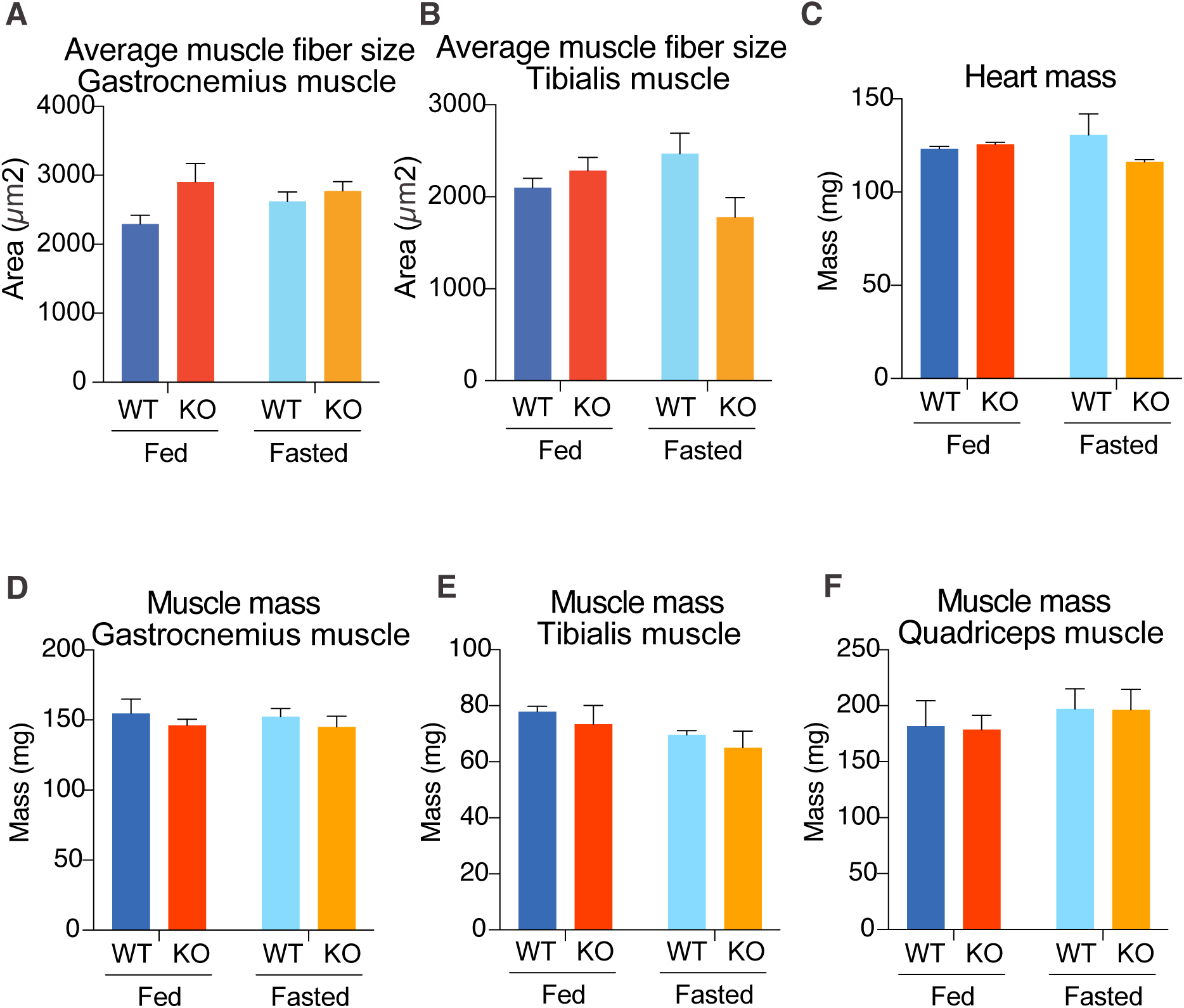
Ism1 ablation does not reduce muscle mass or muscle fiber size in all tissue locations. (A-B) Quantification of average muscle fiber area in gastrocnemius (A) and tibialis (B) in a blinded fashion by two independent investigators (WT fed, *n* = 3; Ism1-KO fed, *n* = 3, WT fasted, *n* = 3; Ism1-KO fasted, *n* = 3, two-way ANOVA, *p < 0.05, **p < 0.01, ***p < 0.001). (C) Heart mass of WT and Ism1-KO mice in the fed or fasted groups (WT fed, *n* = 3; Ism1-KO fed, *n* = 3; WT fasted, *n* = 3; Ism1-KO fasted, *n* = 3, two-way ANOVA). (D-F) Muscle masses of gastrocnemius (D), tibialis (E), and quadriceps (F) of WT and Ism1-KO mice in the fed or fasted groups (WT fed, *n* = 3; Ism1-KO fed, *n* = 3; WT fasted, *n* = 3; Ism1-KO fasted, *n* = 3, two-way ANOVA). Bar graphs show mean ± SEM.

**Figure 4—source data 1**

Raw data related to Figure 4B, C, and E-G.

**Figure 5—source data 1**

Raw data related to Figure 5A-D and F-H.

**Figure 5—source data 2**

Uncropped western blot images with relevant bands labelled.

**Figure 6—source data 1**

Raw data related to Figure 6A and C-G.

**Figure 6—source data 2**

Uncropped western blot images with relevant bands labelled.

## Data availability

All data generated or analyzed during this study are included in the manuscript and supporting files. The phosphoproteomics dataset has been deposited to ProteomeXchange Consortium through JPost PXD031719 (JPST001484)(Okuda et al., 2017). The code for all analysis related to phosphoproteomic data is available at https://github.com/Svensson-Lab/Isthmin-1/tree/F442A_phosphoproteomics. The single-cell RNA sequencing data was re-analyzed from a previously published dataset (Baht et al., 2020). All generated reagents will be shared upon request, including Ism1 recombinant proteins and Ism1-KO and Ism1-flox mice. Further information and requests for resources and reagents should be directed to and will be fulfilled by the Lead Contact, Dr. Katrin J. Svensson.

